# Precision cell-state silencing of injured afferents alleviates spontaneous neuropathic pain

**DOI:** 10.64898/2026.10.08.757642

**Authors:** Andrew H Cooper, Paschalina Chrysostomidou, Ana Jurado Santos, Hening Cui, Siobhan Adesida, Magdalena Redondo Canales, Alice Bakina, Allison M Barry, Georgios Baskozos, David L Bennett, Greg A Weir

**Author notes:** These authors contributed equally to the work (Andrew H Cooper, Paschalina Chrysostomidou).

## Abstract

Excessive primary afferent activity sustains neuropathic pain, but safe and effective therapeutic inhibition has proved elusive. Realising the promise of gene therapies to meet this challenge requires strategies to identify and selectively target the neuronal populations responsible. Here, we use chemo- and optogenetic approaches to demonstrate that activity in injured afferents drives spontaneous pain but is dispensable for normal sensory function, thereby creating a therapeutic opportunity. Guided by chromatin accessibility profiling, we create an injury-responsive viral vector that confers >95% specificity for injured sensory neurons. Combined with the translationally relevant chemogenetic PSAM^4^-GlyR, this targeting module effectively silences hyperexcitable human sensory neurons and alleviates spontaneous neuropathic pain in mice. These findings establish precision cell - state targeting of sensory neurons as a promising therapeutic strategy for chronic pain.

## Introduction

Neuropathic pain arises because of a lesion or disease of the somatosensory nervous system. It is common, affecting 7-10% of adults, and prevalence is rising because of increasing rates of diseases that result in nerve damage, including diabetes and cancer (*1*). Treatment approaches are not keeping pace with this escalating clinical need, and most patients are failed by current medications due to both poor efficacy and tolerability, and live with often disabling pain (*2*).

A major clinical complaint of patients suffering from neuropathic pain following peripheral nerve injury is spontaneous pain i.e., pain occurring in the absence of a stimulus. This pain often co-exists alongside sensory loss and sensory hypersensitivity (allodynia or hyperalgesia). Primary afferent hyperexcitability is considered a key driver of painful symptoms, as demonstrated by the effectiveness of targeted local anaesthetic nerve block to relieve both spontaneous pain and sensory hypersensitivities in patients (*3*). However, from a translational standpoint, unbiased pharmacological silencing of primary afferents risks the loss of protective sensory function, in addition to off-target motor, CNS and cardiac side effects. Given the rapidly expanding repertoire of molecular tools for on-demand neuronal silencing, gene therapy offers a potentially selective approach. For this potential to be realised, the neuronal populations driving pathological pain need to be identified an d approaches developed to restrict transgene expression to just these neurons.

Partial injury to peripheral nerves commonly results in neuropathic pain, and excessive activity occurs in primary afferents directly injured by the insult (*4*), as well as intact neighbouring afferents (*5*); raising the question of how these two populations contribute to the pain experienced. This has been a key question in the field for decades (*6–8*) with important implications for targeting strategies yet is so far unresolved. A further challenge is that in most clinical neuropathic pain disorders injured and uninjured afferents are closely intermingled within sensory circuits, precluding selective anatomical targeting.

Here, we have used Atf3^CreERT2^ mice (*9*) to target chemo- and optogenetic silencers to just injured neurons following experimental nerve injury. We demonstrate that selective silencing of injured afferents is sufficient to reduce spontaneous pain but does not impact stimulus-evoked mechanical or thermal sensitivity. This creates a therapeutic opportunity, where spontaneous pain could be selectively alleviated, while sparing normal somatosensation. To advance this aim, we have used single-nucleus ATAC-sequencing to profile naïve and injured sensory neuron open chromatin accessibility and have identified a DNA regulatory element active in injured but not naïve sensory neurons. Incorporated into viral vectors, this DNA sequence is sufficient to drive high levels of transgene expression in injured sensory neurons, with virtually no expression in naïve counterparts. Finally, we use this regulatory element in combination with a chemogenetic actuator, PSAM^4^-GlyR, to demonstrate the clinical translatability of this gene therapy approach, which has efficacy across species, in both rodent and human models of nerve injury.

## Results

### Injured afferents drive spontaneous pain

Previous lesioning studies produced conflicting data regarding the relative contribution of activity in directly injured and adjacent intact afferents to neuropathic pain (*10*, *11*). We devised a genetic strategy to selectively silence injured afferents in an experimental nerve injury model and thus determine their contribution to pain behaviours. We induced expression of a modified glutamate-gated Cl^-^ channel (GluCl), consisting of α and β-subunits, which is activated exclusively by ivermectin (IVM) to produce prolonged peripheral neuron silencing (*12*). GluCl expression was restricted to injured afferents via intrathecal (i.t.) viral delivery of Cre-dependent constructs (*13*) into Atf3^CreERT2^ mice, in which Cre is expressed *de novo* in sensory neurons following axonal injury (*15*)(Atf3-GluClαβ mice) (**Fig. 1A**). The functional GluCl channel requires αβ heterodimers and so mice expressing only the β subunit (Atf3-GluClβ) were used as control. Animals subsequently underwent spared nerve injury (SNI), which results in a mixture of axotomized and intact neurons in the lumbar DRGs. Using this approach, GluCl was expressed by injured DRG neurons, with negligible expression in intact adjacent afferents (**Fig. 1B and Fig. S1A and B**). Efficient and selective expression was evident regardless of tamoxifen dosing regimen (**Fig. S1b)**, and so subsequent experiments used a dose (3x 30mg/kg) previously shown to induce efficient recombination with minimal axotomy-independent labelling (*14*). An advantage of the slow off kinetics of IVM is that silencing effectiveness can be assessed *ex vivo* following *in vivo* IVM dosing (*12*). GluClαβ^+^ DRG neurons derived from IVM-treated Atf3-GluClαβ animals exhibited a decreased input resistance and concomitant reduced excitability when compared to neurons isolated from control mice (**Fig. S1C-F**), confirming effective GluCl-mediated silencing (*12*).

**Fig. 1.**
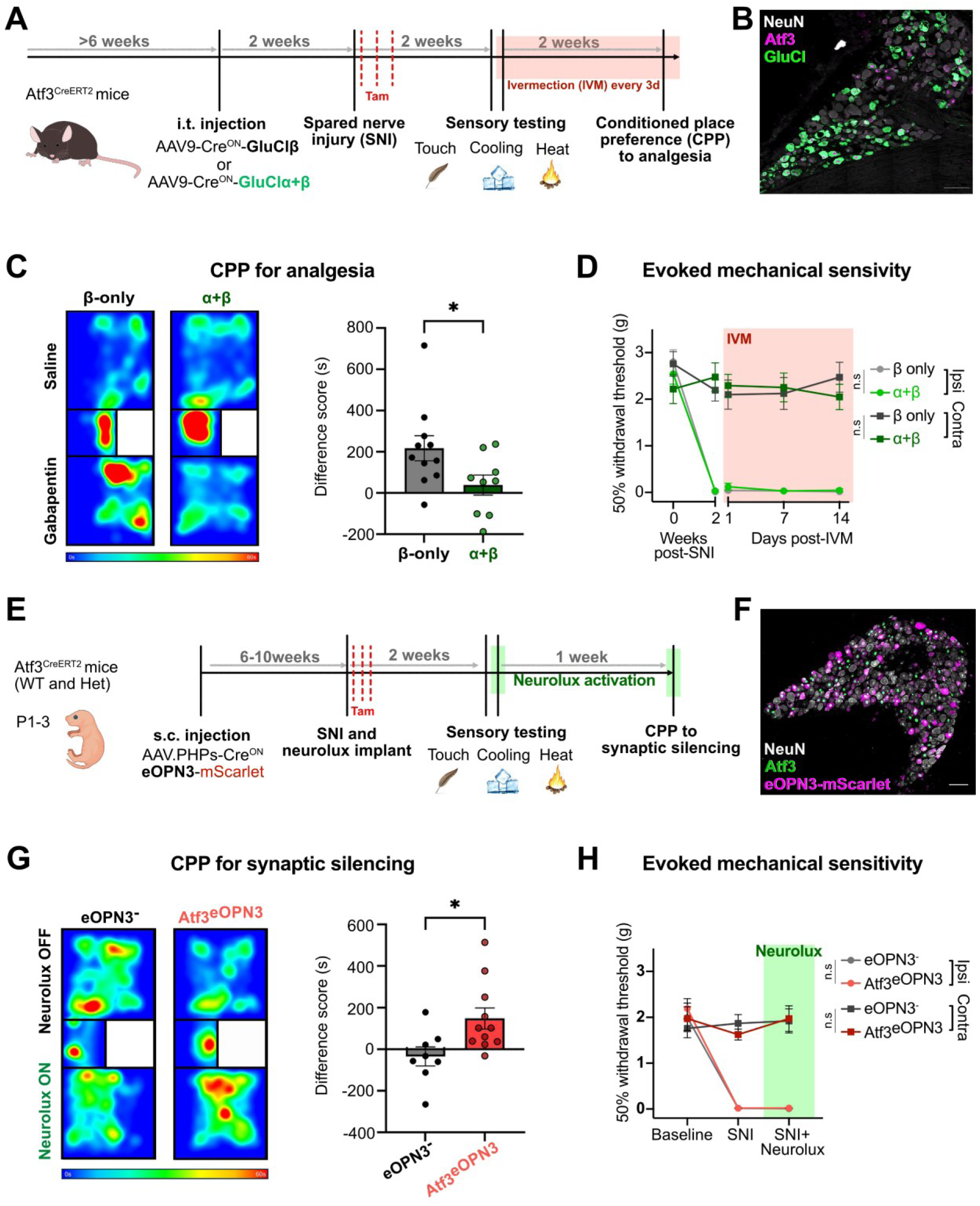
Activity in injured afferents drives spontaneous pain following nerve injury. (**A**) Schematic showing design of experiment to silence injured sensory neurons after SNI using Cre^ON^-GluCl activated by ivermectin (IVM). (**B**) L4 ipsilateral DRG two weeks after SNI, from an Atf3-GluClαβ animal. (**C**) *Left*, heatmap of relative time spent by a control (β-only) and an experimental (α+β) mouse in each chamber following conditioning during CPP testing. *Right,* Reduced CPP to gabapentin (100 mg/kg) in Atf3-GluClαβ animals (unpaired t-test; *P* = 0.039, *n* = 9-11). (**D**) Longitudinal assessment of mechanical sensitivity showing no effect of injured afferent silencing ipsilateral or contralateral to nerve injury. 3-way RM ANOVA, Time x Injury x GluCl interaction: *P* = 0.49; *n* = 8. (**E**) Schematic showing design of experiment to prevent neurotransmission from injured sensory neuron central terminals after SNI using eOPN3 activated by a spinal µLED (Neurolux) implant. (**F**) L4 ipsilateral DRG four weeks after SNI, from an ATF3^CreERT2^ animal injected with Cre^ON^-eOPN3-mScarlet. (**G**) *Left*, heatmap of relative time spent by a control (wild-type) and an experimental (Atf3^CreERT2+/-^) mouse in each chamber following conditioning to µLED activation. *Right,* µLED activation elicits preference in eOPN3 expressing mice but not control littermates (unpaired t-test; *P* = 0.0196, *n* = 8-11). (**H**) Longitudinal assessment of mechanical sensitivity showing no effect of silencing injured afferent neurotransmission ipsilateral or contralateral to nerve injury. 3-way RM ANOVA, Time x Injury x Neurolux *P* = 0.73; *n* = 6-8. Scale bars represent 100µm.

Peripheral nerve injury produces spontaneous pain with a prominent negative affective component, which can be assayed in rodents by conditioned place preference (CPP) to analgesia (*15*). We tested whether chronic IVM-mediated silencing of injured afferents was sufficient to alter signs of spontaneous pain. Gabapentin induced CPP in control mice following nerve injury, consistent with relief of spontaneous pain (**Fig. 1C and Fig. S1G and H**). In contrast, gabapentin failed to induce CPP in Atf3-GluClαβ mice during IVM-mediated silencing, indicating that selective inhibition of injured DRG neurons attenuates spontaneous pain, despite preservation of signalling in adjacent intact afferents. Crucially, IVM administration did not alter evoked mechanical, cold, or heat sensitivity in the contralateral hindpaw or stimulus-evoked hypersensitivity in the adjacent intact sural dermatome (**Fig. 1D and Fig. S1I–K**), indicating that silencing does not disrupt sensory processing conducted through uninjured pathways.

We used a second means of silencing to reproduce and extend our results. In contrast to GluCl-mediated membrane shunting (*12*), the Gi-coupled opsin eOPN3 inhibits presynaptic neurotransmitter release (*16*) and allowed us to directly test whether acute silencing of injured afferent neurotransmission produces CPP (**Fig. 1E**). eOPN3 expression was selectively targeted to injured neurons, with minimal expression in adjacent intact DRG neurons, and no expression in the ipsilateral DRG of wildtype littermates (**Fig. 1F and Fig. S2A and B**). Given the lack of eOPN3 expression in animals lacking cre, we interpret the low levels of channel expression in Atf3^-^ neurons of the ipsilateral DRG to be likely due to Atf3 being down - regulated in axotomized afferents by the time of analysis (four weeks post-SNI) (*17*), as opposed to cre-independent eOPN3 expression. Optogenetic inhibition of injured sensory afferent terminals induced CPP, while µLED activation had no effect in wildtype littermates that lacked eOPN3 expression (**Fig. 1G and Fig. S2C and D).** As observed with GluCl-mediated inhibition, selective suppression of injured afferent transmission had no effect on mechanical, cold, or heat hypersensitivity, nor did it alter baseline sensory function (**Fig. 1H and Fig. S2E-G**).

Collectively, these findings demonstrate that selective silencing of injured afferents alleviates spontaneous pain while preserving normal sensory function, thus making these neurons an attractive analgesic target.

### Injured afferents display a unique open chromatin accessibility landscape

There is to date no means to selectively target damaged neurons without genetically engineered animals. Incorporating cis-regulatory elements (CREs) into engineered AAV has proven highly effective in driving cell-type selective expression in the CNS (*18*). Given the major gene expression changes that occur in sensory neurons following damage (*19*), we predicted that identifying injury-activated CREs may allow us to engineer AAVs to selectively target injured sensory neurons. To do so, we profiled chromatin accessibility of injured and intact DRG neurons using Avil^FlpO^;Atf3^CreERT2^;RC::FLTG transgenic mice, which facilitate differential and high-fidelity fluorescent labelling of both populations (**Fig. 2A and B**) (*14*). Fluorescently labelled neuronal populations were isolated by FACS from DRG of both naïve and nerve-injured mice four weeks following SNI (**Fig. S3A-C**). Incorrect gating in some cell collections resulted in the inclusion of non-neuronal cells into each sample, but this did not result in cross contamination of either neuronal type (**Fig. S3D**). We performed snATAC-seq on samples, which following quality controls (**Fig. S4A-C**) and removal of non-neuronal cells (**Fig. S4D**), resulted in a dataset of 1,248 naïve, 3,146 contralateral, 2,901 ipsilateral intact and 617 ipsilateral injured neurons (**Fig. S4E**). Given their concordance, we integrated naïve and contralateral samples into a single naïve atlas and performed unsupervised clustering based on gene-activity scores, which resolved nine clusters. We annotated clusters using previously described marker genes (*20*) and could identify most known sub-populations, albeit with a likely under-representation of some large diameter afferent populations (**Fig. 2C**). To corroborate these annotations, we transferred cell-type labels from a reference scRNA-seq atlas of DRG neurons (*20*) which supported our assignments and resolved several subtypes within our broader clusters (**Fig. 2D**). Following joint analysis of naïve and injured (GFP^+^) neurons, GFP^+^ cells were found to largely segregate from naïve sub-populations and form distinct clusters (**Fig. 2E**). To discover potentially useful CREs, we separately compared injured neurons to our two control groups (contralateral and naïve) and identified 2,853 significantly enriched consensus peaks (**Fig. 2F**). Peaks were predominantly located in distal intergenic and intronic regions and showed most enrichment for Activator Protein-1 (AP-1) family motifs, which are known regulators of the transcriptional response to injury (*19*) (**Fig. 2G**). We prioritised 132 *de novo* injury-induced peaks whose nearest gene is upregulated across multiple sub-populations of injured sensory neurons (*21*). One enriched peak overlapped with the transcriptional start site (TSS) of the regeneration-associated gene, Sprr1a (*22*) (**Fig. 2H**), and was notable for three reasons: (1) Sprr1a gene expression is induced up to 60-fold following nerve injury, across neurons of all sizes (*22*, *23*), and has very limited basal nervous system expression (*24*, *25*), (2) the peak was virtually absent from all naïve sub-populations, and (3) we noted that the upstream region of this peak included many predicted binding sites of the core injury-induced transcription factors Atf3, Jun and Stat2 (**Fig. 2I**) (*19*). These observations led us to hypothesise that there may be sufficient regulatory sequences proximal to the core Sprr1a promoter to render this region useful as an injury-responsive regulatory element (*26*).

**Fig. 2.**
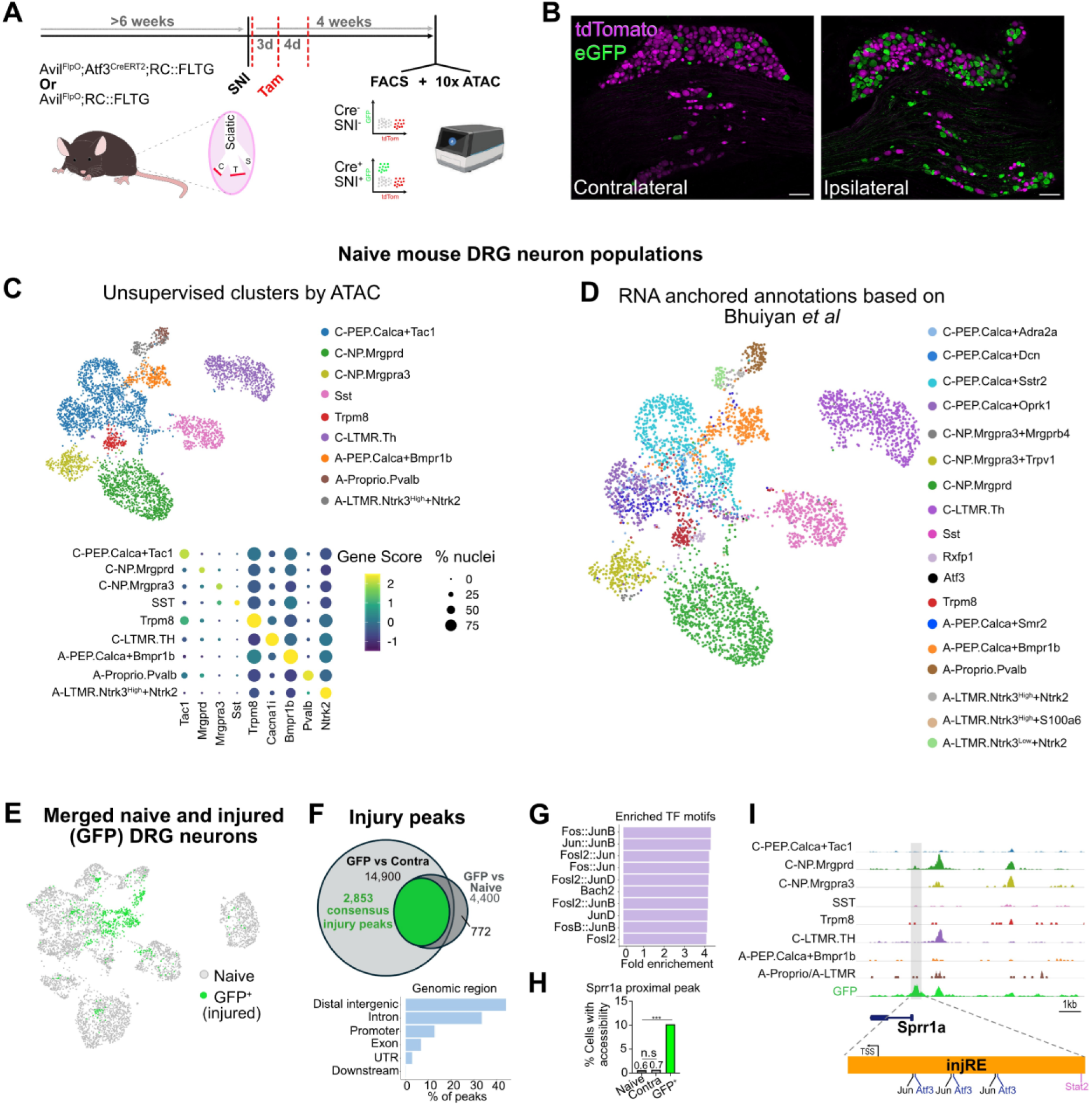
Injured afferents have differential open chromatin accessibility to naïve sensory neuron sub-populations. (**A**) Experimental timeline for the isolation of naïve and injured sensory neurons. (**B**) L4 ipsilateral and contralateral DRG four weeks after SNI from an Avil^FlpO^;Atf3^CreERT2^;RC::FLTG mouse, in which injured afferents are labelled with eGFP and naïve neurons with tdTomato. Scale bar represents 100µm. (**C**) *Top*, UMAP projection of 4,394 neurons integrated from naïve and contralateral samples. *Bottom*, Dot plot displaying ATAC gene activity scores for canonical sub-population marker genes used to manually annotate clusters. (**D**) Subtype annotation by reference-based label transfer from Bhuiyan et al.(*20*). (**E**) UMAP projection of 617 injured neurons merged with naïve atlas. (**F**) Significantly differential peaks from independent GFP *vs* contralateral and GFP *vs* naïve comparisons, and their genomic locations. Of the 3,633 consensus peaks, 2,853 were more accessible in injured neurons. Note, greater contralateral sample size led to a higher number of peaks identified from this comparison. (**G**) Top 10 enriched TF motifs across 2,853 significantly enriched consensus peaks. (**H**) Percentage cells with accessibility at the peak proximal to Sprr1a (log₂FC 3.78, FDR 1.3×10⁻¹⁶; pairwise Fisher’s exact tests between groups). Note, the modest absolute fraction in injured cells is consistent with the sparsity of single-cell ATAC data (the median cell has non-zero signal at ∼9% of peaks). (**I**) Normalised chromatin accessibility at the genomic loci of Sprr1a across injured and naïve sub-populations. Location of injRE1 in relation to Sprr1a and ATAC peak is illustrated, as are JASPAR-predicted (*57*) transcription factor binding sites for Jun, Atf3 and Stat2.

### Injury-responsive element allows for selective targeting

To functionally test this CRE, we cloned a 500bp region centred on the middle of the ATAC peak (designated injRE1) into a GFP reporter construct. This region included the core promoter region of Sprr1a, which negated the need for a minimal promoter as is often required for enhancer AAVs. We packaged the reporter construct into an engineered capsid that preferentially transduces neurons of the peripheral nervous system (*27*) and delivered AAV.PHP.s.injRE1.GFP systemically to neonatal mice (**Fig. 3A**). One week following nerve injury in adulthood, 49.7 ± 7.1% of Atf3^+^ (axotomized) neurons in ipsilateral DRG neurons were GFP labelled (**Fig. 3B**) yet only 0.2 ± 0.2% of contralateral DRG neurons were also (**Fig. 3C**). Of note, some contralateral labelled neurons were Atf3^+^ and may represent a distinct molecular state, or a low level of injury in naïve DRG (*28*). Population distributions of neuron size showed no difference between GFP^+^ and reference contralateral DRG neurons (**Fig. 3D**). Central terminal labelling confirmed the stark unilateral labelling, which was also evident across laminae in the ipsilateral dorsal horn (**Fig 3E**), consistent with broad GFP expression across sub-populations of axotomized sensory neurons.

**Fig. 3.**
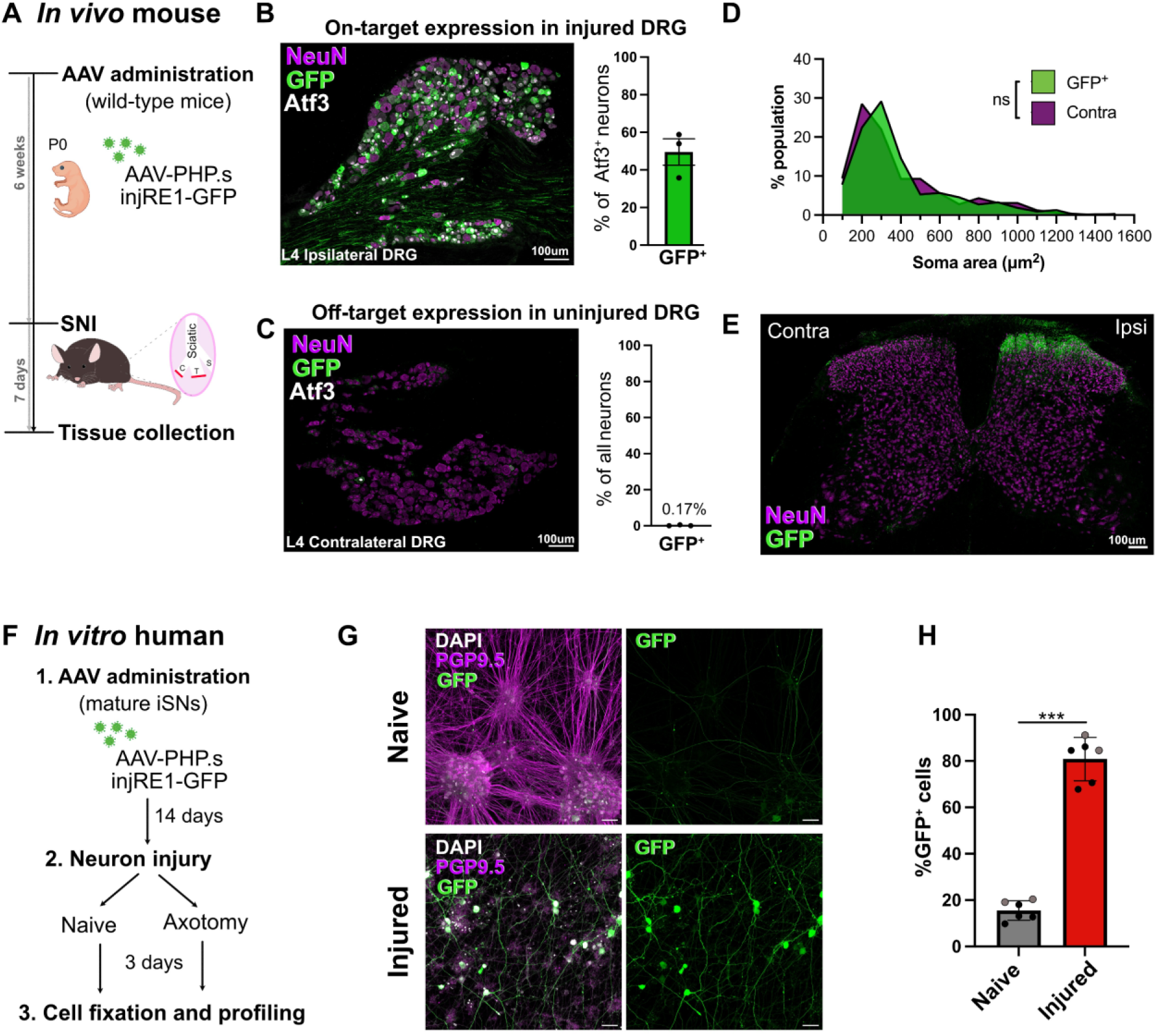
Injury-response module directs high and selective reporter expression in injured afferents. (**A**) Experimental design for testing injRE1 *in vivo*. (**B**) On-target GFP expression in injured (ipsilateral Atf3^+^) DRG neurons and, (**C**) off-target GFP expression in contralateral DRG neurons. *n* = 3 mice for data shown in (B) and (C). (**D**) Population distribution of labelled and reference (contralateral) DRG neurons by cross-sectional area. Kolmogorov–Smirnov tests of cumulative distributions; *n* = 323-550 cells pooled from three mice. (**E**) labelling of GFP sensory neuron central terminals in regions innervated by injured nerve but not uninjured territories. (**F**) Schematic of *in vitro* experiment to test injRE1 in a human sensory neuron injury paradigm. (**G**) Example images of naïve and injured iSNs expressing GFP, driven by injRE1. (**H**) Percentage of iSNs expressing GFP. *n* = 6, from 2 independent iPSC lines: black circles - SBAD2, grey circles - SBAD3. Unpaired t-test *** *P* <0.001 Scale bars represent 30µm.

The upstream region of Sprr1a shares ∼60% sequence similarity between mouse and human, and so we tested whether our mouse derived injRE1 sequence could also drive high and selective expression in injured human sensory neurons. Induced pluripotent stem cell-derived sensory neurons (iSNs) exhibit molecular and physiological features of endogenous sensory neurons and initiate an injury response upon axotomy (*29*). AAV.PHP.s.injRE1.GFP treatment of iSNs resulted in sparse and low level GFP expression under naïve conditions that was markedly increased upon axotomy (**Fig. 3F-H**).

Together, these experiments confirm that injRE1 drives high transgene expression selectively in injured over intact sensory neurons across species and therefore represents a valuable injury-responsive module for viral targeting.

### injRE1-PSAM^4^-GlyR as a therapeutic approach

Chemogenetic methods for controlled neuromodulation have been developed for discovery neuroscience to interrogate pain circuits, but their translational potential is also being increasingly recognised (*30*). Critical to their successful translation to the clinic will be selective expression in only the intended neuronal types. We tested whether the injRE1 module could be combined with ion channels/receptors to functionally silence injured sensory neurons and alleviate pain.

We first delivered DNA constructs containing inward-rectifying potassium channel Kir2.1 downstream of injRE1 into dissociated (and therefore injured) mouse DRG neurons (**Fig. 4A and B**). This channel produces a potassium leak current that hyperpolarises the membrane potential and has previously been used to reduce sensory neuron hyperexcitability (*31*). Expression driven by injRE1 was sufficient to induce large inward-rectifying currents (**Fig. 4C and D**) and a marked hyperpolarisation of the membrane potential compared to control neurons (**Fig. 4E**). These altered membrane properties rendered Kir2.1 expressing cells hypoexcitable to depolarising stimuli (**Fig. 4F**).

**Fig. 4.**
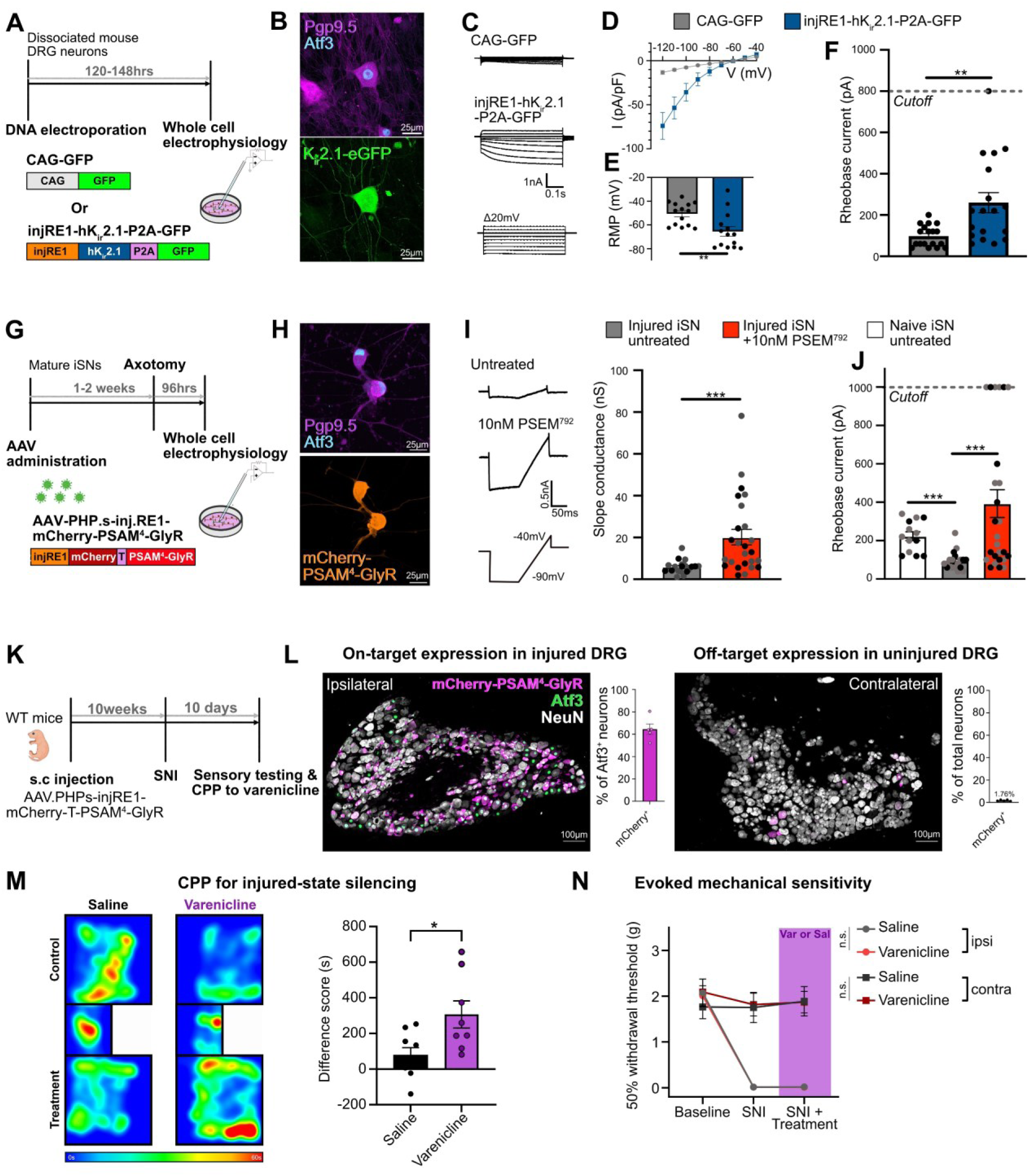
On-demand silencing of injured afferents alleviates spontaneous pain behaviour. (**A**) Testing of injRE1 combined with the inwardly rectifying potassium channel, hKir2.1. (**B**) A mouse DRG neuron expressing hKir2.1 six days after dissociation (and therefore injury). Scale bars = 25 µm. (**C**) Voltage clamp traces from cells electroporated with control (CAG-GFP) or injRE1-hKir2.1-P2A-eGFP during depolarizing voltage steps from -120mV to -40mV. (**D**) Current-voltage relationship derived from (C). hKir2.1 expressing cells show a clear inward rectifying current at hyperpolarised potentials, indicating expression of functional channel. (**E**) Experimental neurons also exhibit a hyperpolarised resting membrane potential (RMP) and require an increased current injection to fire an action potential, (**F**) compared to control. DRG neurons that failed to respond to an 800pA stimulus were given this value. Data in (B-F) derived from two mice. Mann-Whitney test, *\*\* P* < 0.01. (**G**) injRE1 combined with mCherry-T-PSAM^4^-GlyR and packaged into AAV. (**H**) AAV-treated iSNs express mCherry following injury. Scale bars = 25 µm. (**I**) uPSEM^792^ elicits an increased slope conductance in injured iSNs expressing mCherry-T-PSAM^4^-GlyR (Mann-Whitney test, *\*\*\* P* < 0.001). *Left*, representative voltage clamp traces and, *right*, quantification. (**J**) Injured iSNs have reduced rheobase compared to naïve counterparts, which is reversed upon uPSEM^792^ treatment. iSNs that failed to respond to 1nA stimulus were given this value. One-way ANOVA followed by Dunn’s multiple comparisons test *\*\*\* P* <0.001. Data in (G-J) derived from 2 independent iPSC lines: *black circles - SBAD2, grey circles - SBAD3*. (**K**) *In vivo* testing of AAV.PHP.s.injRE1.mCherry-T-PSAM^4^-GlyR. (**L**) Example images and quantification of mCherry-T-PSAM^4^-GlyR expression in ipsilateral and contralateral DRG neurons two weeks post-SNI. Scale bars = 100 µm. (**M**) *Left*, heatmap of time spent by AAV-treated animals in each chamber following conditioning. *Right,* varenicline elicits a preference in AAV-treated mice following nerve injury, whereas saline control does not (unpaired t-test; *P* = 0.016, *n* = 8-9). (**N**) Longitudinal assessment of mechanical sensitivity showing no effect of varenicline ipsilateral or contralateral to nerve injury (3-way RM ANOVA; Time x Injury x Treatment: *P* = 0.632, *n* = 8-9).

Constitutive ion channel expression may have translational drawbacks and so we next assessed whether chemogenetic actuators were as amenable to combination with injRE1. PSAM^4^-GlyR is an effector with the advantage of temporal control and reversibility, which has previously been shown effective in acute and chronic pain models (*32*, *33*). We therefore generated AAVs containing injRE1 and mCherry-PSAM^4^-GlyR and tested the effectiveness of the approach in our humanised *in vitro* model of nerve injury (**Fig. 4G**). iSNs robustly expressed mCherry-PSAM^4^-GlyR following injury (**Fig. 4H**), and neurons treated with the ultrapotent agonist, uPSEM^792^, displayed a greater background conductance compared to untreated counterparts (**Fig. 4I**), consistent with functional channel activation (*32*). As previously reported, axotomised iSNs had lowered rheobase compared to naïve counterparts (*29*) (**Fig. 4J**). However, this hyperexcitability was reversed upon treatment with uPSEM^792^ (**Fig. 4J**). These *in vitro* data demonstrate that injRE1 drives sufficient expression of actuators to effectively silence injured mouse and human neurons.

Given the translational potential of selective PSAM^4^-GlyR expression, we next sought to assess efficacy of injRE1.mCherry-PSAM^4^-GlyR for reversing pain behaviour *in vivo*. Neonate wild-type mice were administered systemically with AAV before undergoing SNI in adulthood (**Fig. 4K**). injRE1 directed high levels of expression in injured afferents (64.5 ± 4.6% of ipsilateral Atf3^+^ DRG neurons) with minimal off-target expression in intact DRG neurons (1.8 ± 0.3% of contralateral DRG neurons) (**Fig. 4L**). Varenicline is an FDA-approved drug and an agonist of PSAM^4^-GlyR that in a control experiment we first demonstrated does not induce CPP in wild-type mice following SNI (**Fig. S5A-C**). However, varenicline was sufficient to induce CPP in AAV.PHP.s.injRE1.mCherry-PSAM^4^-GlyR treated animals following nerve injury (**Fig. 4M and Fig. S5D-E)**. As with our earlier studies relying on transgenic targeting, selective varenicline-mediated silencing of injured afferents did not impact normal mechanosensation (**Fig. 4N**). These data support the translational potential of a combined injury-responsive module and chemogenetic silencer to treat the affective, spontaneous component of neuropathic pain.

## Discussion

We show that activity in directly injured primary afferents drives spontaneous pain behaviours but does not contribute to normal sensory processing, making them an attractive cellular locus for analgesic drug targeting. Exploiting the major injury-evoked changes in gene regulation, we designed an approach to precisely target these neurons with a silencing ‘effector’. We generated the first single nucleus chromatin accessibility dataset of labelled injured sensory neurons and used this to identify an injury-responsive cis-regulatory element (CRE) that confers highly selective transgene expression in sensory neurons only following injury. Combining this element with inhibitory effectors we demonstrate the ability to reduce the electrical activity of injured rodent and human sensory neurons, and reverse spontaneous pain behaviours while sparing normal somatosensation in a neuropathic pain model. These data illustrate the feasibility of targeting sensory neurons based on cell state and advance the concept of developing cell selective gene therapies for chronic pain.

Studies of primary afferent types that contribute to pain behaviours after nerve injury have mostly focussed on neuronal populations defined by naive molecular profile (*34–36*). Here, we have instead focussed on cell state and in doing so have revisited a long-standing debate on the relative contributions of activity in injured and adjacent intact afferents to pain behaviours (*6*, *37*). Our results support a fundamental role for directly injured afferents in driving spontaneous pain but not in mediating evoked hypersensitivities. This has critical implications for analgesic design, highlighting a need to focus on molecular targets expressed on injured afferents. While our viral targeting approach does not achieve 100% targeting, and therefore we cannot exclude a limited overlap, the substantial impacts that we observed on spontaneous pain behaviour mean that we can clearly dissociate the contribution of injured afferents to evoked hypersensitivity and spontaneous neuropathic pain behaviour.

Recent efforts have generated single-cell chromatin accessibility atlases for naïve rodent and human sensory neurons, which have, and will continue to be, highly valuable for targeting of naïve-state molecular sub-populations (*38–40*). Ours is the first study to profile the epigenome of purely injured sensory neurons and here we have prioritised accessible regions based on proximity to a set of genes dysregulated following nerve injury. In addition to painful symptoms, loss of sensation is a major clinical problem in many forms of neuropathic pain. Therefore, an important rationale for selectively targeting injured sensory afferents is to leave the protective roles of spared uninjured afferents intact to prevent injury (for instance painless burns and ulceration) and joint deformity (such as Charcot joints). Larger screens may identify further genomic regions to investigate, for example, that offer more selective targeting of sub-populations of injured neurons. For instance, a recent study by Zeng *et al*, defined a role for a sub-population of injured C-fibre afferents marked by expression of Sstr2, in mediating a novel behavioural readout of paroxysmal pain (*41*).

We have shown that pathological neuronal states can themselves provide the regulatory specificity needed to target and therapeutically silence the neurons driving disease while sparing their healthy counterparts. The injury response of sensory neurons, including the *de novo* up-regulation of Sprr1a, shares overlap with the response of other neuronal types to injury and degeneration (*42*, *43*). The approach may therefore have broader use in neurological contexts where disease is driven by neuronal damage and where gene therapy strategies are being developed, such as Parkinson’s, Amyotrophic Lateral Sclerosis and traumatic brain and spinal cord injury.

Distal enhancer sequences are usually favoured to provide high levels of selective expression in AAV vectors, without the need for the larger sequences common with endogenous promoters. injRE1 overlaps with the core promoter of Sprr1a yet an only 500bp sequence is sufficient to drive high levels of expression with remarkable selectivity; *in vivo* specificity ranged from 95-99%, which is as high as the best performing enhancer sequences defined in large CNS-wide screens (*44*, *45*). A key determinant on the translatability of designer AAVs will not only be their specificity for a target but also that they drive therapeutically meaningful transgene levels. In this regard, injRE1 provided sufficient expression of a number of different inhibitory effectors to successfully reduce excitability of neurons *in vitro,* and represents the first CRE shown to be effective at targeting injured neurons and reversing a disease phenotype *in vivo*.

In summary, we identify directly injured sensory neurons as a key therapeutic target for chronic neuropathic pain and establish injury-responsive cell-state targeting as a strategy for their selective therapeutic silencing.

## Acknowledgments

We thank John Craig and Iain Plenderleith for technical assistance genotyping transgenic mouse lines. Schematics included in Figure 4 include images that were adapted from images provided by Servier Medical Art, licensed under CC BY 4.0. For the purpose of open access, the author has applied a CC BY public copyright licence to any Author Accepted Manuscript (AAM) version arising from this submission.

## Author contributions

Conceptualisation: AHC, PC, DLB, GAW; Methodology: AHC, PC, HC, DLB, GAW Formal analysis: AHC, PC, HC, AMB, GAW; Investigation: AHC, PC, AJS, SA, MRC, AB, GAW; Writing-original draft: AHC, PC, GAW; Writing-review and editing: AHC, PC, AJS, HC, SA, MRC, AB, AMB, GB, DLB, GAW; Visualisation; AHC, PC, HC, GAW Supervision; GB, DLB, GW; Funding acquisition: DLB, GB, GAW

## Data, code, and materials availability

The sequencing data generated during this study will be deposited in the NCBI Gene Expression Omnibus (GEO) and made available upon publication. All other data and resources will be made available upon reasonable request to the corresponding author.

## Materials and Methods

### Animals

Animal procedures were performed under a UK Home Office Project Licence and in accordance with the UK Home Office (Scientific Procedures) Act (1986). All studies were approved by the Animal Welfare and Ethical Review Body of the University of Glasgow and conform to the ARRIVE guidelines. Mice were group-housed in humidity and temperature-controlled rooms with access to food and water *ad libitum*, on a 12-hour light-dark cycle, and with environmental enrichment. Mice were handled and habituated to testing equipment for 30 min/day for three days prior to all behavioural testing, and all procedures took place during the light cycle. All experimental cohorts included both male and female mice and were born in our facility. Mice were randomly assigned to groups using a random number generator before experiments began.

The following mouse lines were used: C57BL/6J (Charles River, UK); Atf3 ^CreERT2^ (Atf3^tm1.1(cre/ERT2)Msra^, a gift from Dr Franziska Denk) (*9*); Avil^FlpO^ (Avil^tm1(flpo)Ddg^, a gift from Prof David Ginty)(*47*); RC::FLTG (B6.Cg-*Gt(ROSA)26Sor^tm1.3(CAG-tdTomato,-EGFP)Pjen^*/J; The Jackson Laboratory RRID:IMSR_JAX:026932)(*48*).

### Plasmid cloning and AAV production

DNA plasmids and AAVs used in this study are detailed in Table S1. The sequence for injRE1 was commercially synthesised and subsequently cloned into a plasmid containing the inverted terminal repeats of AAV2 flanking injRE1, eGFP, woodchuck hepatitis virus post-transcriptional regulatory element (WPRE), and a human growth hormone polyadenylation signal, by BrainCase (Wuhan, China). The eGFP sequence was replaced by that of mCherry-T-PSAM4-GlyR or hKir2.1.P2A.eGFP commercially by VectorBuilder, (Chicago, USA) to generate pAAV.injRE1.mCherry-T-PSAM4-GlyR and pAAV.injRE1.Kir2.1.P2A.eGFP, respectively. Cre^On^-GluCl AAV was packaged into AAV9 and injRE1.mCherry-T-PSAM4-GlyR into AAV.PHP.s by the Viral Vector Facility (University of Zurich, Switzerland). injRE1.eGFP was packaged into AAV.PHP.s by BrainCase (Wuhan, China). Other AAVs used were purchased from repository stocks of the Viral Vector Facility (University of Zurich, Switzerland).

### Systemic viral vector and drug delivery

#### Neonatal AAV injections

AAVs packaged with a PHP.s serotype were injected neat into neonatal mice (0-2 days old) via subcutaneous (s.c.) injection (10 µL) at the nape of the neck.

#### Tamoxifen injections

Tamoxifen was administered via an intraperitoneal (i.p.) injection of 20 mg/mL tamoxifen (Sigma Aldrich) dissolved in corn oil, either 3 times (0, 3-4 and 7 days post-SNI) at 30-50 mg/kg (Atf3^CreERT2^), or once at 7.5 mg/kg on the day of SNI.

#### Systemic administration of pharmacological agents

Gabapentin (100 mg/kg; Sigma, UK) or varenicline (0.3 mg/kg; Tocris, UK) were dissolved in 0.9% saline and administered via i.p. injection into the lower right or left quadrant of the abdomen using a 32G insulin syringe. 1% Ivermectin (Noromectin, Norbrook Laboratories Ltd, UK) was administered via an i.p. injection at 0.5 µL/g of mouse’s weight once every 3 days.

### Surgical procedures

#### Intrathecal cannulation

Intrathecal delivery of viral vectors in 6 to 12-week-old mice was performed as previously described (*12*) and took place two weeks prior to SNI surgery. Anaesthesia was induced with 5% isoflurane and maintained at 1% to 2% as required. Mice were placed on a stereotaxic frame and after surgical exposure, the body of the T11 lumbar vertebra was clamped. The dura was exposed by careful dissection of soft tissue between T10 and T11 vertebrae and punctured with a 30 G needle, before a small cannula (polytetrafluoroethylene tubing, 0.008” outer diameter × 0.004” internal diameter, Braintree Scientific) was inserted 11 mm caudally into the subarachnoid space. Eight microlitres of AAV (4 μL of neat AAV9-Cre^ON^-GluClβ plus 4 μL of neat AAV9-Cre^ON^-GluClα, or phosphate-buffered saline) was then infused at a rate of 1 μL/min. Following a 1-min rest period to limit backflow, the cannula was slowly removed. Mice received post-operative analgesia (carprofen (10 mg/kg) and buprenorphine (0.1 mg/kg)) for 3 days after surgery.

#### Spared nerve injury surgery

Spared nerve injury (transection of the common peroneal and tibial branches of the sciatic nerve; SNI) was performed as described previously (*49*). Anaesthesia was induced with 5% isoflurane and maintained at 1% to 2% as required. Analgesia, consisting of carprofen (10 mg/kg) and buprenorphine (0.1 mg/kg), was provided perioperatively. The left hindpaw was secured with tape in hip abduction, and the operative field (lateral surface of the thigh) was shaved. Ophthalmic ointment was applied to the eyes, and the shaved area was swabbed with chlorhexidine solution. A longitudinal incision was made in the skin at the lateral mid-thigh. Using blunt dissection, an opening was made between the biceps femoris and semitendinosus exposing the sciatic nerve and its peripheral branches (sural, tibial, and common peroneal nerves). The common peroneal and tibial nerves were ligated using 6-0 Vicryl suture (Ethicon, USA) and a 1-2 mm piece distal to the suture was removed using spring scissors. The muscle was then closed with one suture (6-0 Vicryl), and the skin incision closed with one 10 mm wound clip (Alzet, USA) or one to two buried horizontal mattress sutures (6-0 Vicryl). Exclusion criteria included autotomy (mutilation of 2 or more toenails or any living tissue in the denervated region), or a lack of development of mechanical hypersensitivity, however no mice were excluded for these reasons.

#### Wireless µLED (NeuroLux) implantation surgery

Surgical implantation of wireless 530 nm µLED spinal implants(*50*) (NeuroLux Inc., IL, USA) took place one week after SNI and one week before beginning further behavioural testing. Anaesthesia and analgesia were provided as described for i.t. cannulation, and mice were placed on a stereotaxic frame. The bodies of the T12 and L1 lumbar vertebra were exposed and clamped. The T13-L1 intervertebral space was exposed, and the L1 spinal process removed with a dental burr. The tip of the implant, gently bent to conform to the curvature of the spinal column, was inserted rostrally, so the µLED was positioned above the dura over the L4 spinal segment. The probe was then glued into position with Histoacryl adhesive (B. Braun Medical Ltd, UK), and the wound closed with buried horizontal mattress sutures (6-0 Vicryl).

### Behavioural assays

All behavioural measurements were performed by an investigator blinded to experimental conditions. Sample sizes were chosen using power calculations based on our own previous studies, using G*Power 3.1.9.7 (α < 0.05 and Power > 0.8). Where responses to multiple evoked pain modalities were assessed, testing was typically performed in a single session in the following order: von Frey, brush, acetone, and plantar radiant heat.

### Conditioned place preference

To test for the affective, spontaneous component of neuropathic pain, we used a three-day conditioning protocol to test for conditioned place preference (CPP) as previously described (*51*, *52*). The custom-built CPP testing apparatus consisted of two side chambers (170 × 150 mm) connected by a centre chamber (70 × 75 mm), 180 mm tall and with infrared-transparent plastic lids. Mice were able to discriminate between the two side chambers based on visual (vertical or horizontal black and white stripes on the walls) and sensory (rough or smooth textured floors) cues. When testing preference to gabapentin or varenicline, a biased chamber assignment was used, i.e., vehicle (saline) was paired to the preferred chamber, and the drug paired to the less-preferred chamber. When testing preference to the activation of a Neurolux implant, chambers were predetermined before the start of the experiment, with half of the apparatuses wired for activation in the left chamber, half for the right-hand chamber. On day 0 (acclimation), mice had free access to explore all chambers for 30 minutes. On days 1 and 2 (preconditioning), mice were again allowed to freely explore for 30 min whilst their position was recorded using an infrared camera and AnyMaze software (Stoelting, Ireland). For conditioning (days 3 to 5), mice received control treatment (i.p. saline for pharmacological conditioning, no intervention for photostimulation conditioning) and were returned to their home cage for 5 min, then confined to their designated control chamber for 30 min. Three to four hours later, mice received the intervention treatment (i.p. gabapentin or varenicline), returned to their home cage for 5 min, and were then placed in the other side chamber for 30 min, or received µLED photostimulation for the full duration. On test day (day 6), mice could freely explore all chambers whilst their position was recorded, as during pre-conditioning, for 30 min. Difference scores were calculated as the time spent in each chamber on test day minus the mean time spent during pre-conditioning. Exclusion criteria included mice spending >90% or <5% of time in a side chamber or >60% of time in the centre chamber during pre-conditioning. 3/23 mice were excluded from Atf3-GluCl experiments, 1/20 from Atf3-eOPN3 experiments, 1/18 for injRE1-PSAM4-GlyR experiments, and no mice were excluded from other CPP experiments. CPP was selected for optogenetic experiments rather than a real-time preference assay due to the inactivation kinetics of eOPN3 (seconds to minutes) (*16*) which would presumably prevent the spatiotemporal association of chamber to pain relief.

### Von Frey assessment of static mechanical allodynia

50% mechanical thresholds at the hindpaw ipsilateral and contralateral to SNI were measured using the von Frey up-down method (*53*). Mice were placed in acrylic boxes, transparent on external sides, but opaque where required to block view of other mice, on an elevated wire mesh platform, for at least 15 min prior to assessment. The stimulus (von Frey applicators, Stoelting, UK) was applied to the hindpaw at the lateral aspect of the glabrous skin, corresponding to the intact sural dermatome, avoiding the dennervated regions that were previously innervated by the tibial or common peroneal nerves. Each paw was stimulated using a predefined set of 8 monofilaments (0.008 to 6 g), starting each trial with an intermediate filament (0.16 g), applied perpendicular to the skin, causing a slight bending, for 3 s. In case of a positive response (rapid withdrawal or licking of the paw within 3 s of removing of the filament, but ignoring normal ambulation or rearing), the next smallest filament was tested, with at least 2 min between stimulations. In case of a negative response, the next larger filament was tested. Each trial continued until 4 measurements beyond the first change in response (i.e., no response then response, or vice versa) were taken. 50% mechanical withdrawal thresholds were calculated using the Dixon up-down statistical method.

### Brush assay of dynamic mechanical allodynia

Dynamic mechanical allodynia was assessed using the brush method. A fine, 5/0 paintbrush was trimmed to a blunt tip and used for stimulation. Mice were placed in acrylic boxes atop a mesh grid as for von Frey testing. The lateral aspect of the glabrous hindpaw skin was gently stroked from heel to toe at a consistent velocity (∼2 cm/s), and scored using the following: (0) a typical response in a naïve animal, consisting of lifting the paw away from the stimulus for less than 1 s; (1) a single rapid, stereotyped withdrawal response or flinch, or sustained lifting of the paw for >1 s; (2) A lift of the paw laterally, above the level of the hip for >1s, or a startled jump response; (3) A stronger stereotyped withdrawal response consisting of withdrawing and flicking the affected limb, and/or licking the affected paw. Four applications were performed per timepoint, with at least 2 min between each, and the mean response score for each mouse was calculated.

### Acetone assay of cold allodynia

Cold allodynia was assessed using the acetone assay. Mice were placed in the same acrylic boxes atop wire mesh as for the von Frey assay. A 200 µL pipette tip was melted to create a flat surface (∼6 mm diameter, ∼45° relative to the longitudinal axis) while maintaining an open lumen and was attached to a 20 µL pipettor. A 10 µL drop of acetone was formed on the flat pipette tip surface and carefully applied to the lateral aspect of the glabrous hindpaw skin without the tip contacting skin. The duration for which the mouse raised or licked the stimulated paw, or exhibited orbital tightening, was recorded, with a cut-off of 30 s. Three measurements were averaged per time point, with at least 5 min between acetone applications.

### Plantar radiant heat assay (Hargreaves test)

Sensitivity to a heat stimulus was performed using a plantar radiant heat apparatus (IITC Inc., USA). Mice were acclimated for at least 15 mins in acrylic boxes on an elevated glass platform maintained at 30°C. A radiant heat source was then aimed through the glass floor at the lateral edge of the hindpaw using the guide light, and then the stimulus light was activated. Latency to paw withdrawal was recorded, ignoring normal ambulation. Cut-off was set at 20 s to prevent tissue damage. Three measurements were averaged per time point, with at least 5 min between stimulations.

### Light stimulation

For behavioural experiments involving opsin activation, testing chambers were wired externally and connected to tuning and power distribution boxes controlled via NeuroLux software, enabling delivery of the appropriate RF signal to activate wireless µLED implants. The stimulation protocol consisted of 1 Hz cycling with 50% on/50% off at 10 W RF power (adapted from (*16*)). For CPP experiments, stimulation was maintained throughout the conditioning session (30 min). For evoked behavioural assays, light stimulation was maintained for the duration of testing each modality, with a 10 min stimulation-free interval between assays.

### Immunohistochemistry, image acquisition and analysis

Mice were terminally anaesthetised with pentobarbital (20 mg) and transcardially perfused with 4% formaldehyde in PBS. L3 to L5 DRG and L4 segments of spinal cords were removed and post-fixed for a further 2 hours, cryoprotected in 30% sucrose overnight, and then embedded in optimal cutting temperature media (OCT; Tissue Tek, USA) or agar. Cells were fixed by 4% paraformaldehyde for 10 minutes at room temperature. Tissue was sectioned on a Leica CM1950 cryostat, with 12-30 µm sections collected serially on Superfrost Plus slides (Epridia, UK) (DRG) or 60 µm sections collected free-floating in PBS (spinal cord). Samples (tissue or cells) were washed with PBS, before being incubated with appropriate primary antibodies (Table S2) diluted in 5% normal donkey serum (NDS) and 0.3% Triton X-100 in PBS for 3 days at 4°C. After PBS washes, samples were incubated with appropriate fluorophore-conjugated secondary antibodies in PBS/NDS/Triton-X100 solution overnight at room temperature. Spinal cord sections were then mounted on Superfrost Plus slides. Slides were washed and coverslipped with VectaShield Vibrance Hardset mounting media (Vector Labs, CA, USA). Samples were imaged using a Zeiss LSM900 Airyscan confocal microscope equipped with 405, 488, 561 and 640 nm diode lasers. Full thickness, tiled, confocal image stacks with a 1-3 µm interval in the Z-axis were obtained through a 20x dry or 40x oil lens with the confocal aperture set to 1 Airy unit or less. All image capture was performed using Zen Blue Edition software (Carl Zeiss Microscopy GmbH, Germany), and analyses were performed using Zen Blue or FIJI(*54*). Representative images consist of orthogonal projections of all captured optical sections.

### snATAC Sequencing

#### Sample collection and Fluorescence activated cell sorting (FACS)

DRG were collected from adult Avil^FlpO^;Atf3^CreERT2^;RC::FLTG mice 4 weeks after SNI and naïve Avil^FlpO^;RC::FLTG mice. Animals were dosed with 50mg/kg tamoxifen via intraperitoneal injection on days 0, 3, and 7 after SNI. Both male and female animals were used in experiments. Ipsilateral and contralateral L3-L5 DRG were extracted following transcardial perfusion with ice-cold PBS. In total, collections were performed from 16 SNI (6 male, 10 female) and 10 naive (no injury) (7 male, 3 female) mice. DRG were enzymatically digested in HBSS containing Collagenase type II (4 mg/ml) and Dispase type II (4.6 mg/ml) at 37°C for 90 minutes. The cells were then washed with mature Neurobasal medium and mechanically dissociated using glass pipettes of two different sizes. Single cell suspensions were passed through a 70µm reversible strainer and samples were transported on ice to the Flow Core Facility at the University of Glasgow for FACS.

Sorting was performed using a BD FACSAria III. A sorter with a 100µm nozzle was used to sort fluorescent cells into tubes containing 100µl Neurobasal media under pressure at 10 psi. Samples were stained with Draq7 dye to exclude dead cells and sorting criteria monitoring the distribution of tdTomato and/or GFP signals and forward and side scatter were selected. Following sorting, cells were centrifuged for 5 minutes at 600 x *g*, resuspended in 200µl of Cryostor media and stored in -150°C until library preparation.

#### Nuclear isolation and ATAC sequencing

Nuclei were isolated from frozen samples according to the protocol provided by 10X Genomics. Cells were washed three times with pre-warmed PBS supplemented with 0.04% bovine serum albumin (BSA) and centrifuged at 500 x *g* for 10 minutes at 4°C. After the last wash, the supernatant was removed, leaving 5µl in the tube. Samples were then incubated with 2X chilled lysis buffer (20mM Tris-HCl, 20mM NaCl, 6mM MgCl2, 0.2% Tween-20, 0.2% IGEPAL Ca-360, 0.02% Digitonin, 2% BSA) for 10 minutes on ice, gently pipetting every 3 minutes. The cells were washed with wash buffer (10mM Tris-HCl, 10mM NaCl, 3mM MgCl2, 1% BSA, 0.1% Tween-20) and centrifuged at 500 x *g* for 15 minutes at 4°C. Nuclei were resuspended in chilled diluted nuclei buffer, centrifuged, and resuspended in nuclei buffer, followed by counting to determine nuclei concentration. Four snATAC-seq libraries were obtained using 10X Genomics Chromium Single Cell ATAC v2.1 chemistry; injured (GFP), ipsilateral (tdTom), contralateral, and naive (refers to age-matched Atf3^CreERT2^ negative, Avil^FlpO^ positive (heterozygous) and RC::FLTG positive (heterozygous) animals which did not undergo SNI). Sequencing was performed on the Illumina HiSeq 4000 System using the 10X Genomics pipeline to a minimum of 50,000 reads per nucleus at the Glasgow Polyomics Facility.

### snATAC-seq analysis

#### Pre-processing, clustering, visualisation, and annotation

Initial pre-processing was performed by the Glasgow Polyomics facility using CellRanger-atac (v2.2) and peaks were called using its default MACS2-based peak calling algorithm. Raw reads were aligned to the mouse reference genome (GRCm38/mm10) and further analysis was carried out using the Signac (version 1.17.1) and Seurat (version 5.5.1) packages in R (version 4.4.1) as previously described(*55*).

Each library was subset to exclude nuclei with peak region fragment counts below the 2nd percentile and above the 98th percentile, percentage reads in peaks and transcription start site (TSS) enrichment below the 2nd percentile and nucleosome signal and blacklist ratio above the 98th percentile. Samples then underwent normalisation and dimensionality reduction using latent semantic indexing (LSI). Non - linear dimension reduction was calculated based on the top 20 principal components (excluding the first component) using the RunUMAP() function while clustering was performed using the FindNeighbors() and FindClusters(algorithm=3) functions.

Following clustering, a pseudo gene expression activity matrix was computed using the GeneActivityMatrix() function. Nuclei with low *Rbfox3* and high *Prx*, *Fabp7*, or *Mpz* gene activity were classed as non-neuronal and were removed from all downstream analyses. Clusters which were found to be double positive for *Rbfox3* and *Prx*, *Mpz*, or *Fabp7* gene activity were also excluded from analyses. Differential gene activity analysis was performed using FindAllMarkers() in Seurat, comparing nuclei in one cluster with all other nuclei. Genes with log2FC > 0.5 and FDR < 0.05 were considered enriched in a given cluster. Clusters were annotated based on the gene activity of known neuronal subtype markers as determined both by differential gene activity analysis and manual inspection of activity using functions such as FeaturePlot() and VlnPlot().

#### Dataset merging and integration

For the integration of the naive and Contra datasets, a unified set of peaks was created and quantified in each dataset using FeatureMatrix(). Following normalisation and linear dimension reduction, the two datasets were merged and then integrated. Briefly, integration anchors between the datasets were identified using reciprocal LSI (FindIntegrationAnchors(reduction= “rlsi”)) and used to correct the LSI embeddings with IntegrateEmbeddings(). This was followed by further non - linear dimension reduction and clustering carried out as described above. Differentially accessible peaks were identified using FindAllMarkers() and analyses looked for peaks with minimum expression in at least 10% of cells (min.pct = 0.1), FDR < 0.05, and log2FC > 0.5 and employed a Wilcox rank-sum test (test.use = ‘wilcox’). For the combined analysis of naive (naive and Contra) and injured (GFP) samples, a unified set of peaks was created for all three datasets as described earlier. The datasets were then merged before undergoing analysis as previously described.

#### Annotation using published datasets

The harmonised cross-species DRG atlas publeshed by Bhuiyan et al. (2024)(*20*) was downloaded from harmonized.painseq.com. Following subsetting to keep only cells originating from mouse samples, gene expression values were anchored onto nuclei from our integrated naive dataset using FindTransferAnchors() and subtype labels were transferred with TransferData().

#### CRE identification and prioritising

Differential accessibility was tested with FindMarkers using a logistic-regression framework (test.use = “LR”) with the number of peak fragments per cell (nCount_peaks) as a latent variable to control for sequencing depth, comparing GFP against each control separately (GFP vs Contra; GFP vs naive). Peaks with FDR < 0.05 and |log₂ fold-change| > 0.25 were called significant. Peaks significant against both controls were first taken together, of these, 5 changed in opposite directions between the two comparisons and were excluded, leaving 3,628 injury-specific peaks with a consistent direction of change (2,853 opened, 775 closed in injured neurons). Peaks with zero control accessibility were treated as pseudocount-inflated and were not ranked by fold-change. Consensus peaks were annotated to genomic features and nearest genes with ChIPseeker (v1.46.1) using TxDb.Mmusculus.UCSC.mm10.knownGene and org.Mm.eg.db, with a promoter region of ±3 kb around the TSS. Position frequency matrices for vertebrate transcription factors were obtained from JASPAR2020 (CORE collection) to assess transcription-factor motif enrichment. Motif occurrences were scanned across peaks with motifmatchr against the mm10 genome (BSgenome.Mmusculus.UCSC.mm10), and enrichment in injury-opened peaks was computed with Signac FindMotifs against a GC-content matched background of accessible peaks. To identify regulatory elements that open *de novo* after injury, we selected consensus injury-induced peaks accessible in fewer than 1% of cells in both control groups (naive and contralateral) that were significantly more accessible in injured neurons. Restricting further to peaks whose nearest gene is transcriptionally upregulated in a dataset generated from high depth RNAseq from purified populations of sensory neurons following spared nerve injury (*21*), yielded 132 *de novo* injury-induced elements, which were manually inspected by plotting their coordinates with CoveragePlot().

### Mouse DRG culture

DRG were rapidly excised and enzymatically digested in HBSS containing dispase type II (4.7 mg/ml) plus collagenase type II (4 mg/ml) (Worthington Biochemical) at 37°C for 90 mins. DRG from along the rostral-caudal axis were used except for experiments concerning AAV.Cre^ON^.GluCl (after SNI), wherein only ipsilateral L3, L4 and L5 tissues were harvested. Mechanically dissociated cells were plated onto laminin/poly-D-lysine (BD Biosciences) treated coverslips in complete Neurobasal® medium [Neurobasal® Plus media supplemented with 2% (v/v) B27 Plus, N2 (1% (v/v) and 1% (v/v) antibiotic-antimycotic (ThermoFisher Scientific)]. Mouse nerve growth factor (50 ng/ml; NGF, PeproTech) and 10 ng/ml glial-derived neurotrophic factor (GDNF, PeproTech) were added to the media and cultures used for experiments 24– 72 h later. For cultures generated from animals treated with AAV.Cre^ON^.GluCl, neurons were plated and maintained in DMEM-F12 with 10% foetal calf serum (ThermoFisher Scientific) instead of complete Neurobasal® medium (owing to different concentration of Cl^-^ ions) and used for experiments within 24 h of plating. Electroporation was performed prior to plating using the Amaxa II electroporation system (Lonza Bioscience). After electroporation, cells were immediately plated as described above and used 5-6 days later, when expression of the plasmid DNA was maximal.

### Whole-cell patch clamp electrophysiology

Whole-cell patch clamp recordings were performed at room temperature (22°C) with a Multiclamp 700B amplifier and Digidata 1550 acquisition system (Molecular Devices). Series resistance was compensated 60–80% to reduce voltage errors. Data were low pass filtered at 2 kHz and sampled at 10 kHz and recordings were analysed by Clampfit 10 software (Molecular Devices). Extracellular solution contained (mM): 140 NaCl, 4.7 KCl, 1.2 MgCl2, 2.5 CaCl2, 10 HEPES and 10 glucose; pH was adjusted to 7.3 with NaOH and osmolarity was set to 315 mOsm. For experiments including DRG neurons expressing Kir2.1, intracellular solution contained (mM): 130 K-gluconate, 10 KCl, 2 MgCl2, 2 ATP-Na2, 0.5 GTP-Na, 10 HEPES, and 0.5 EGTA; pH was adjusted to 7.3 with KOH and osmolarity set to 305 mOsm. For experiments including DRG neurons and iSNs expressing GluCl or PSAM^4^-GlyR, intracellular solution contained (mM): 100 K-gluconate, 28 KCl, 1 MgCl2, 5 MgATP, 10 HEPES, and 0.5 EGTA; pH was adjusted to 7.3 with KOH and osmolarity set to 305 mOsm. Intracellular solution was loaded into patch pipettes of 4–8 MΩ tip resistance made from filament borosilicate glass capillaries (1.5 mm outer diameter, 0.84 mm inner diameter; World Precision Instruments). For all recordings, one minute following establishing whole cell access, a one-minute recording was made in bridge mode to derive resting membrane potential. Input resistance (RInput) was derived from the membrane potential deflection caused by a 20pA hyperpolarising current pulse at −60 mV.

### Recordings from Kir2.1 expressing DRG neurons

Resting membrane potential values were corrected during recordings for a liquid junction potential (LJP) of −15mV. To determine the minimal current injection to fire an action potential (rheobase), cells were depolarised from resting membrane potential by current steps (50 ms) of increasing magnitude (Δ20 pA) until an action potential was generated. *I/V* curves were derived from the current elicited by incremental voltage steps (Δ10mV) ranging from −120 to -40 mV, from a resting potential of -60mV.

### Recordings from GluCl or PSAM^4^-GlyR expressing neurons

Resting membrane potential values were not corrected for a LJP of -13.4mV. Rheobase was determined by 20-50ms current steps of 10-100pA increments from a holding potential of -60mV. In voltage clamp, 100ms ramps from −90 mV to −40 mV were used to measure membrane conductance. uPSEM^792^ (Tocris Bioscience) was solubilised in ddH2O to make a 1mM stock solution, which was then further diluted directly in extracellular solution and bath applied during the duration of recordings.

### Stem cell culture and maintenance

Experiments were performed using two previously characterised human induced pluripotent stem cell (iPSC) lines: SBAD2, derived from skin fibroblasts of a 51-year-old male donor, and SBAD3, derived from skin fibroblasts of a 36-year-old female donor. iPSCs were maintained in mTeSR™ Plus medium (STEMCELL Technologies, #100-0276) on Matrigel-coated tissue culture dishes (Corning, #354234) at 37°C in a humidified incubator containing 5% CO₂. Cells were routinely passaged as aggregates using ReLeSR (STEMCELL Technologies, #05872). For differentiation, iPSCs were dissociated into single cells using Accutase (Thermo Fisher Scientific, #A1110501) and seeded onto Matrigel-coated culture plates at a density of 4 × 10^4^ cells/cm² in mTeSR Plus supplemented with CEPT 1000X (Thermo Fisher Scientific, #A56799). Cells were expanded for 24–48 h until they reached approximately 40–50% confluency before sensory neuron differentiation was initiated.

### iPSC-derived sensory neuron differentiation

Human iPSCs were differentiated into induced sensory neurons (iSNs) using a modified version of the protocol described (*56*). Differentiation was initiated (Day 0) by replacing mTeSR Plus medium with KnockOut Serum Replacement (KSR) medium consisting of KnockOut DMEM (Thermo Fisher Scientific, #10829018), 15% KnockOut Serum Replacement (Thermo Fisher Scientific, #10828-028), 1% GlutaMAX (Thermo Fisher Scientific, #35050038), 1% non-essential amino acids (Thermo Fisher Scientific, #11140050), 1% penicillin/streptomycin and 100 μM β-mercaptoethanol (Thermo Fisher Scientific, #31350-010). Neural induction was achieved by dual-SMAD inhibition using SB431542 (10 μM, Tocris, #1614) and LDN-193189 (500 nM, Tocris, #8150) from Days 0–5. Beginning on Day 2, cultures were additionally treated with CHIR99021 (3 μM, Tocris, #4423), SU5402 (10 μM, STEMCELL Technologies, #73914) and DAPT (10 μM, Tocris, #2634) to promote neural crest specification and sensory neuron lineage commitment. The culture medium was gradually transitioned from KSR to N2/B27 medium in 25% increments beginning on Day 4, reaching 100% N2/B27 medium by Day 10. N2/B27 medium consisted of Neurobasal medium (Thermo Fisher Scientific, #21103049) supplemented with 2% B27 minus vitamin A (Thermo Fisher Scientific, #12587010), 1% N2 supplement (Thermo Fisher Scientific, #17502048), 1% GlutaMAX and 1% penicillin/streptomycin. On Day 12, differentiating cultures were dissociated with Accutase (Thermo Fisher Scientific, #A1110501) and replated onto Matrigel-coated glass coverslips at approximately 3 × 10^4^ cells/cm² in N2/B27 medium supplemented with CHIR99021 (3 μM), CEPT and four recombinant neurotrophic factors: BDNF (PeproTech, #450-02), NT-3 (PeproTech, #450-03), NGF (PeproTech, #450-01) and GDNF (PeproTech, #450-10), each at a final concentration of 25 ng/mL. CHIR99021 was included for a maximum of four days following replating.

To minimise contamination by proliferating non-neuronal cells, cytosine β-D-arabinofuranoside (Ara-C; 4 μM, Thermo Fisher Scientific) was applied for 8–24 h between Days 14 and 20, depending on the extent of non - neuronal cell growth. From Day 25 onwards, cultures were maintained in N2/B27 Plus medium comprising Neurobasal Plus medium (Thermo Fisher Scientific, #A3582901), 2% B27 Plus supplement (Thermo Fisher Scientific, #A3582801), 1% N2 supplement and 1% GlutaMAX, supplemented with 25 ng/mL BDNF, GDNF, NGF and NT-3. Fifty percent media changes were performed twice weekly, and neurons were maintained until at least Day 45 before use in experiments.

pAAV.PHP.s.injRE1.GFP and pAAV.PHP.s.injRE1.mCherry-T-PSAM^4^-GlyR were added to cultures at 1ul/500ul of media, resulting in an approximate MOI of 0.4-2×10^5^. Mature iSNs were axotomized as described previously, by manually lifting from their growth plates and dissection into smaller aggregates using a scalpel blade. Cell clusters were mechanically dissociated to single cells using fire-polished glass pipettes of decreasing bore size and subsequently plated onto Matrigel-coated coverslips in N2/B27 Plus medium supplemented with CEPT and growth factors. Experiments including iSNs treated with pAAV.PHP.s.injRE1.mCherry-T-PSAM^4^-GlyR were performed following a refinement to the procedure wherein aggregates were sequentially incubated in 10U/ml papain (Worthington Biochemical) and dispase type II (1.17 mg/ml) plus collagenase type II (1 mg/ml) (Worthington Biochemical) each for 10 minutes at 37 °C. Aggregates were washed in N2/B27 Plus medium with 10% FBS before performing mechanical titration and plating as described.

### Statistical analysis

Data are expressed as mean ± SEM unless otherwise specified, and *P* values of less than 0.05 were considered significant. Statistical analyses were performed in Prism 10 or 11 (GraphPad Software Inc., USA) or R using unpaired or paired t-tests, Mann-Whitney tests, one-, two-or three-way RM ANOVAs, or mixed effect analyses where appropriate. If a main or interaction ANOVA effect was significant, then Šídák’s or Tukey’s multiple comparisons tests were performed. To compare population distributions of soma cross-sectional area or volume, Kolmogorov-Smirnov tests were performed.

