## Supplementary Materials for "Precision cell-state silencing of injured afferents alleviates spontaneous neuropathic pain"

### Supplemental Figures

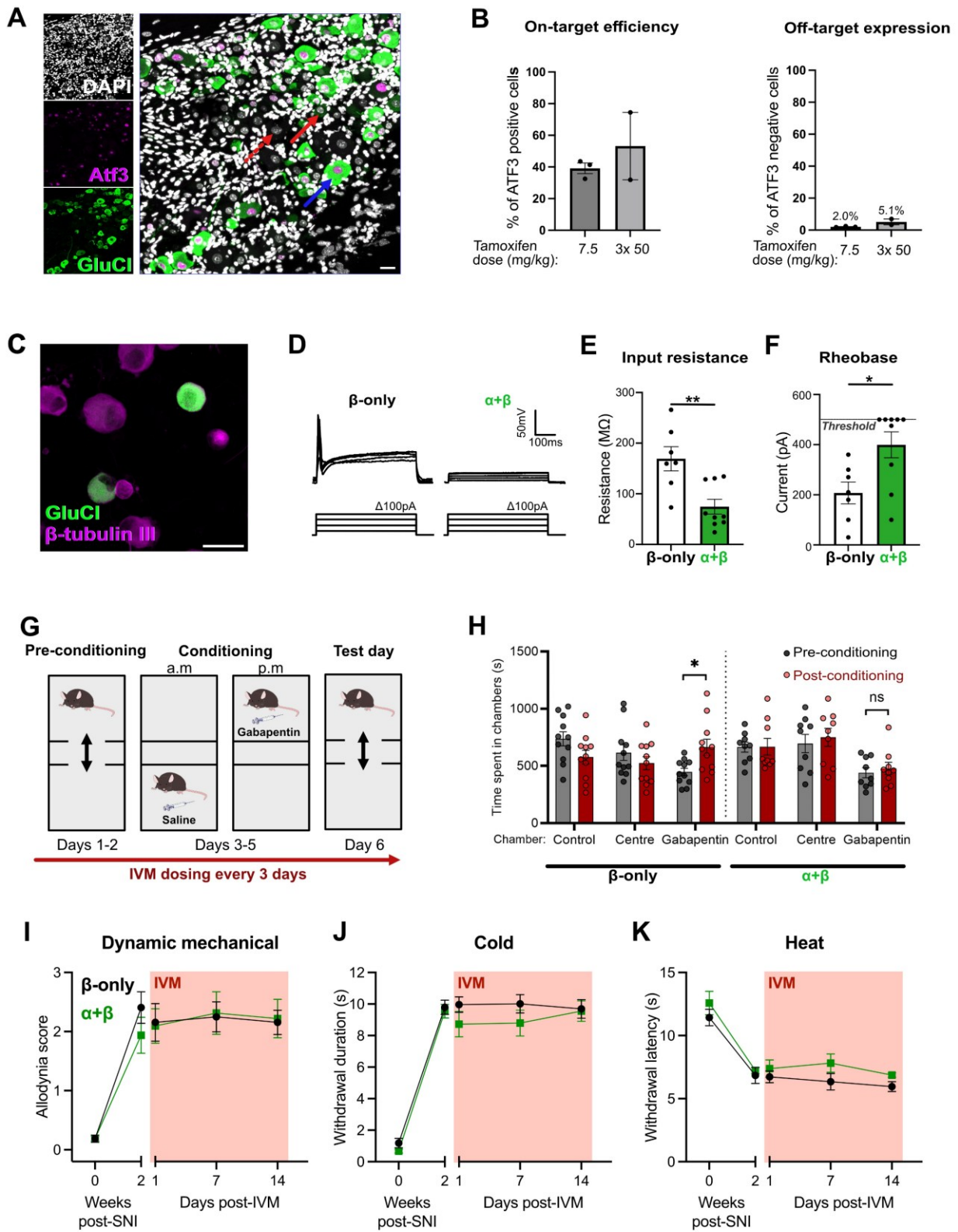

**Fig. S1. Cre<sup>ON</sup>-GluCl targeting of injured afferents.**

(A) Example image of ipsilateral L4 DRG from an Atf3-GluClαβ animal two weeks post-SNI. Red arrows note example Atf3<sup>-</sup> cells with (solid) or without (dashed) GluCl expression and blue arrow notes an Atf3<sup>+</sup>GluCl<sup>+</sup> neuron. Scale bar = 25 μm. (B)

Percentage of Atf3<sup>+</sup> (left) and Atf3<sup>-</sup> (right) neurons that express GluCl following either a single or multiple administrations of tamoxifen. Note that even high tamoxifen dosing produces only negligible off target labelling. **(C)** Mouse DRG neurons dissociated from an Atf3-GluCl$\alpha\beta$ animal post-SNI. Scale bar = 30 μm. **(D)** Representative action potential firing in response to prolonged current (500ms) stimulus of neurons derived from either Atf3-GluCl$\alpha\beta$ or Atf3-GluClβ-only animals post-SNI and dosed with 5mg/kg IVM one day prior to dissociation. **(E)** Input resistance and **(F)** depolarising current required to fire an action potential for these same neurons. Data in (E) and (F) derived from $n = 2$ mice per group. Mann-Whitney test, \*\* $P < 0.01$, \* $P < 0.05$. **(G)** Experimental design for CPP experiments. **(H)** Time spent in chambers pre- and post-conditioning during CPP experiments. Atf3-GluClβ-only mice developed a preference to the gabapentin-paired chamber, in contrast to Atf3-GluCl$\alpha\beta$ mice. Mixed-effects analysis; Chamber x Conditioning x GluCl interaction: $P = 0.196$; Tukey's multiple comparisons: \* $P < 0.05$; $n = 9-11$. **(I-K)** Longitudinal assessment of dynamic mechanical (brush), cold (acetone) and heat (Hargreaves) sensitivity showing no effect of silencing injured afferents ipsilateral or contralateral to nerve injury. 2-way RM ANOVAs; Time x GluCl interactions, Brush: $P = 0.56$, Acetone: $P = 0.66$, Hargreaves: $P = 0.86$; $n = 8$.

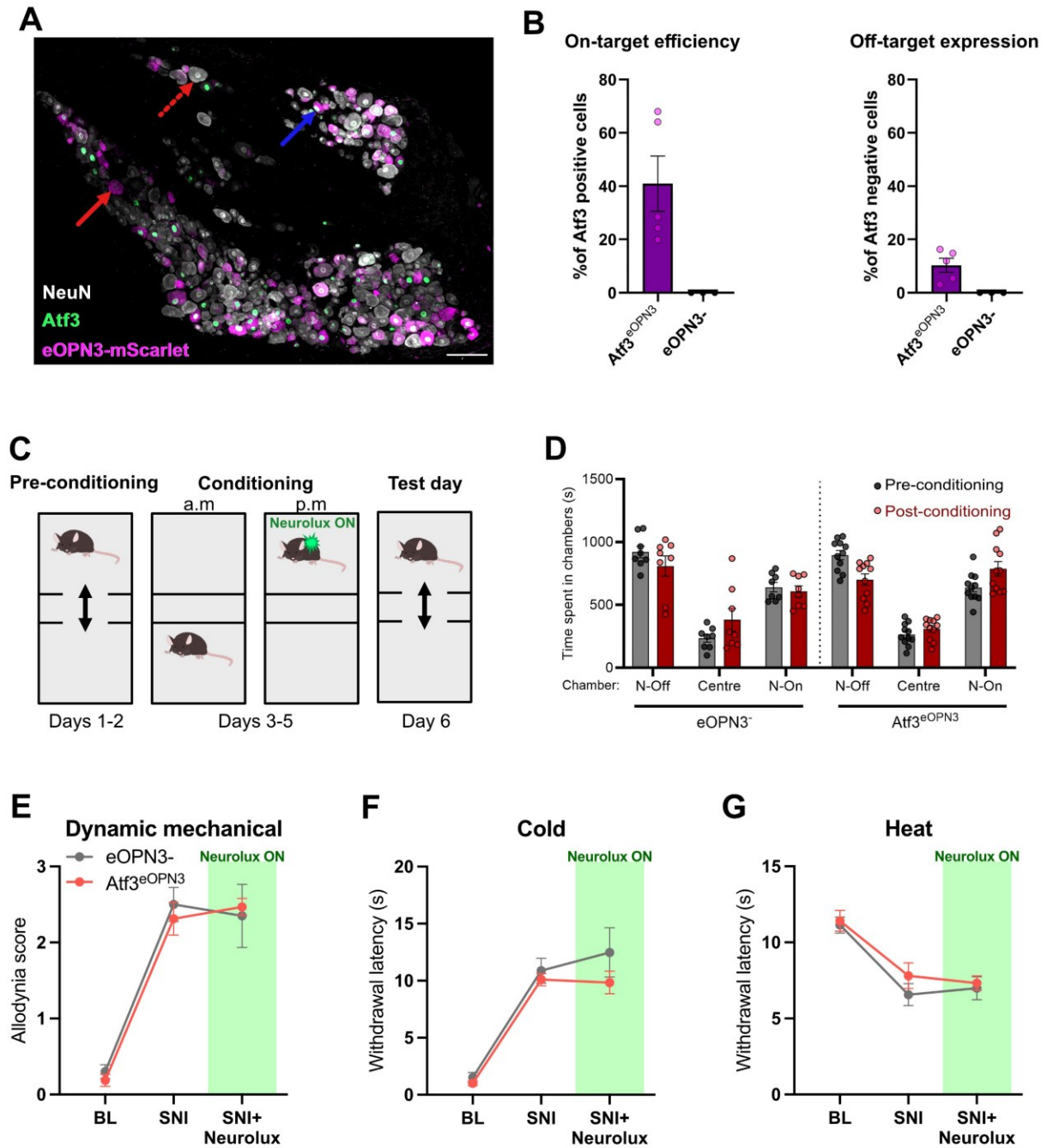

**Fig. S2. CreON-eOPN3-mScarlet targeting of injured afferents.**

(A) Example image of ipsilateral L4 DRG two weeks post-SNI from an $Atf3^{CreERT2}$ animal injected with Cre<sup>ON</sup>-eOPN3-mScarlet ($Atf3^{eOPN3}$). Red arrows note example $Atf3^+$ cells with (solid) or without (dashed) eOPN3 expression and blue arrow notes an $Atf3^+eOPN3^+$ neuron. Scale bar = 100 μm. (B) Percentage of $Atf3^+$ (left) or $Atf3^-$ (right) DRG neurons that express eOPN3 in $Atf3^{CreERT2}$ and wild-type littermates. Data is derived from $n=3-5$ mice per group. (C) Experimental design for CPP experiments. (D) Time spent in chambers paired with μLED stimulation on or off, pre- and post-conditioning, during CPP experiments. Mixed effect analysis; Chamber × Conditioning × eOPN3 interaction: $P = 0.02$; $n = 8-11$. (E-G) Longitudinal assessment of dynamic mechanical (brush), cold (acetone) and thermal heat (Hargreaves) sensitivity showing no effect of silencing injured afferent neurotransmission ipsilateral or contralateral to nerve injury. 2-way RM ANOVAs; Time x eOPN3 interactions, Brush: $P = 0.55$, Acetone: $P = 0.36$, Hargreaves: $P = 0.71$; $n = 6-8$.

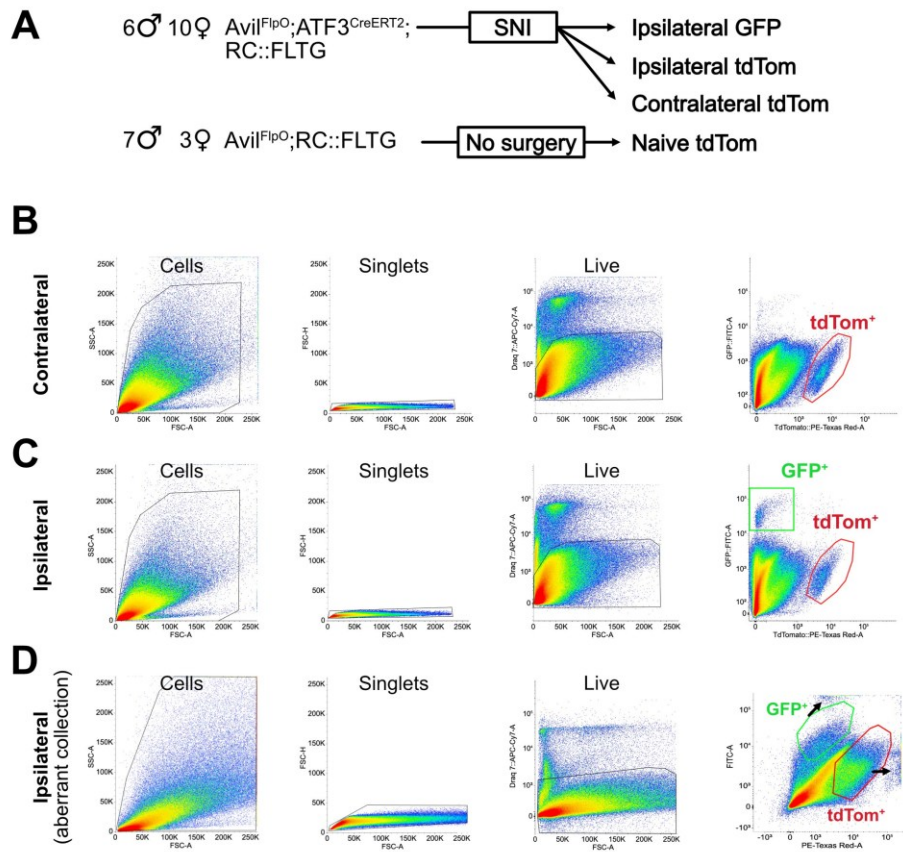

**Fig. S3. Sample information and FACS gating strategy.**

(A) Number and genotype of animals pooled together for each of the described samples (all L3-5 DRG). (B) and (C) Ipsilateral and contralateral samples were gated based on forward (FSC) and side scatter (SSC) to enrich events for cells, single cells based on the forward scatter area (FSC-A) and height (FSC-H), live cells (i.e., those which were Draq7 far-red negative), and collected as either a GFP positive or tdTomato positive populations. (D) Example of an ipsilateral sample where GFP and tdTomato gates were inappropriately placed to capture GFP and tdTomato negative populations. Black arrows highlight GFP positive and tdTomato positive events off the axis scale.

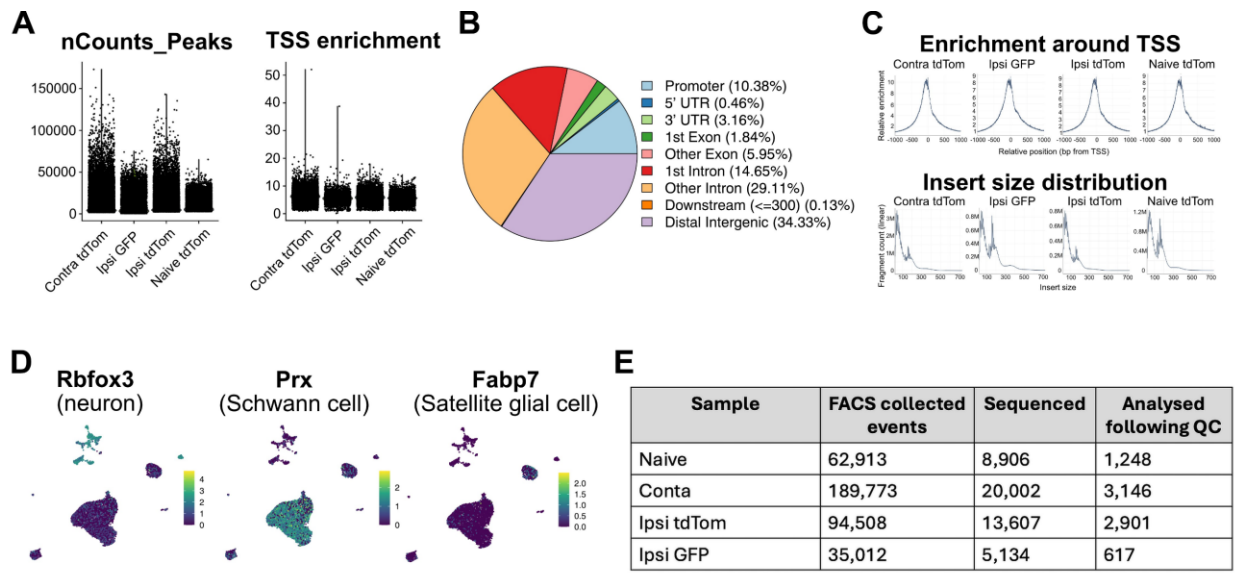

**Fig. S4. Quality control metrics and post-hoc filtering of snATAC samples.**

(A) Total number of fragments per nucleus and transcriptional start site (TSS) enrichment for each of the samples. TSS enrichment measures read density around TSS regions compared to background regions. (B) Proportion of peaks pooled from all samples linked to different genomic annotations., UTR; untranslated region. (C) TSS plot indicating enrichment around the TSS and insert size distribution plot demonstrating the distribution of transposase accessible fragments sequenced. (D) Feature plots of unfiltered naïve sample showing clear distinction between clusters with high Rbfox3 gene activity scores compared to those with high activity scores for non-neuronal cell types such as Schwann cells and satellite glial cells. (E) Sample characteristics, including total cell numbers for each group following quality control filtering steps.

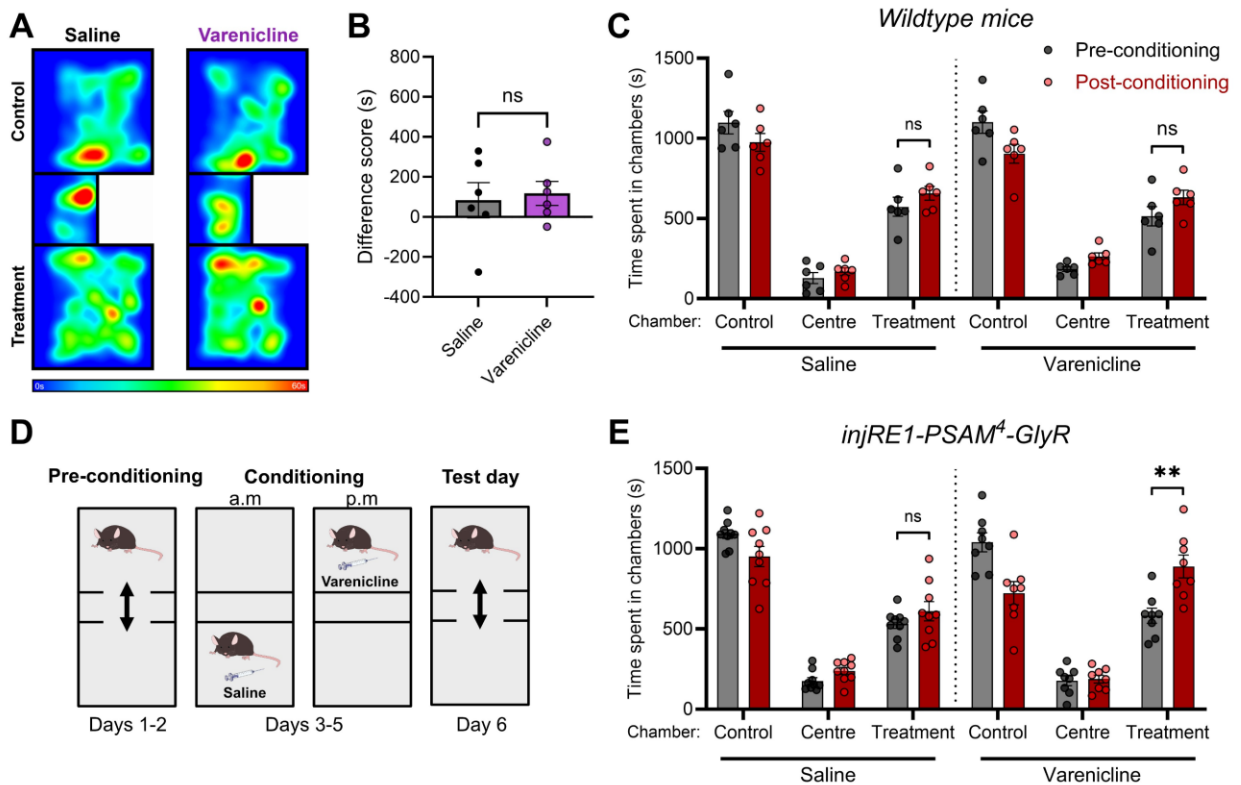

**Fig S5. Varenicline does not induce CPP in the absence of PSAM-GlyR expression**

(A) Heatmap of relative time spent in chambers following conditioning, in mice two weeks after SNI that lack PSAM<sup>4</sup>-GlyR expression. (B) No significant difference was observed in difference scores measured in the treatment-paired chamber between saline- and varenicline-treated mice (unpaired t-test; $P = 0.75$, $n = 6$). (C) No difference was observed in the time spent in chambers pre- and post-conditioning to saline or varenicline (0.3 mg/kg) during CPP testing (Mixed-effects analysis; Chamber x Conditioning x Treatment interaction: $P = 0.64$, $n = 6$). (D) Experimental design for CPP experiments. (E) Time spent in chambers paired with varenicline (0.3 mg/kg) or saline, pre- and post-conditioning, during CPP experiments in wildtype mice treated with AAV.PHP.s.injRE1.mCherry-PSAM<sup>4</sup>-GlyR (Mixed effect analysis; Chamber x Conditioning x Treatment interaction: $P = 0.004$; $n = 8-9$; Tukey's multiple comparisons \*\* $P = 0.0005$).

### Supplemental Tables

**Table S1. Adeno-associated virus used in the study.**

| **Viral Construct** | **Source** | **Titre** | **Identifiers** |
| --- | --- | --- | --- |
| AAV.PHPs-hEF1a-Cre<sup>ON</sup> eOPN3-mScarlet | University of Zurich, Viral Vector Facility | 1.2x10<sup>13</sup>vg/ml | V817 |
| AAV9-sCAG-Cre<sup>ON</sup>-GluClα | University of Zurich, Viral Vector Facility | 1.6x10<sup>13</sup>vg/ml | Custom production |
| AAV9-sCAG-Cre<sup>ON</sup>-GluClβ | University of Zurich, Viral Vector Facility | 1.3x10<sup>13</sup>vg/ml | Custom production |
| AAV.PHPs-injRE1-eGFP | Brain Case, Wuhan, China | 2x10<sup>12</sup>vg/ml | Custom production |
| AAV.PHPs-injRE1-mCherry-T-PSAM<sup>4</sup>-GlyR | University of Zurich, Viral Vector Facility | 1.2x10<sup>13</sup>vg/ml | Custom production |

**Table S2. Primary and secondary antibodies or histological stains used in the study.**

| **Antibody/stain** | **Source** | **Identifiers** | **Working dilution** |
| --- | --- | --- | --- |
| Anti-GFP (Chicken polyclonal) | Abcam plc.,<br>Cambridge, UK | Cat#: ab13970<br>RRID:AB_300798 | 1:1000 |
| Anti-NeuN (Guinea pig polyclonal) | Synaptic Systems,<br>Göttingen, Germany | Cat#: 266004<br>RRID:AB_2619988 | 1:500 |
| Anti-mCherry (Rat monoclonal) | Invitrogen; Thermo<br>Fisher Scientific, UK | Cat#: M11217<br>RRID:AB_2536611 | 1:500 |
| Anti-Atf3 (Rabbit polyclonal) | Novus Biologicals,<br>Minneapolis, MN,<br>USA | Cat#: NBP1-85816<br>RRID:AB_11014863 | 1:500 |
| Anti-PGP9.5 (Chicken polyclonal) | Invitrogen; Thermo<br>Fisher Scientific, UK | Cat#: PA1-10011<br>RRID:AB_1088162 | 1:1000 |
| Anti-β III tubulin (Mouse monoclonal) | Abcam plc.,<br>Cambridge, UK | Cat#: ab78078<br>RRID:AB_2256751 | 1:500 |
| Alexa Fluor 488 Donkey Anti-Chicken IgY (Donkey polyclonal) | Jackson<br>ImmunoResearch,<br>PA, USA | Cat#: 703-545-155<br>RRID:AB_2340375 | 1:500 |
| Alexa Fluor 647 Donkey Anti-Guinea pig IgG (Donkey polyclonal) | Jackson<br>ImmunoResearch,<br>PA,<br>USA | Cat#: 706-605-148<br>RRID:AB_2340476 | 1:250 |
| Alexa Fluor Plus 555 Donkey anti-Rat IgG Highly Cross-adsorbed (Donkey polyclonal) | Invitrogen; Thermo<br>Fisher Scientific, UK | Cat #: A48270<br>RRID:AB_2896336 | 1:500 |
| Alexa Fluor 647 Donkey Anti-Rabbit IgG (Donkey polyclonal) | Jackson<br>ImmunoResearch,<br>PA, USA | Cat# 711-605-152<br>RRID: AB_2492288 | 1:250 |
| Rhodamine Red-X Donkey Anti-Rat IgG (Donkey polyclonal) | Jackson<br>ImmunoResearch,<br>PA, USA | Cat#: 712-295-153<br>RRID:AB_2340676 | 1:100 |
| Streptavidin, Pacific Blue conjugate | Molecular Probes;<br>Thermo Fisher<br>Scientific, UK | Cat#: S11222 | 1:500 |
